# X upregulation is not global and extent of upregulation differs between ancestral and acquired X-linked genes

**DOI:** 10.1101/2021.07.18.452817

**Authors:** C H Naik, K Hari, D Chandel, MK Jolly, S Gayen

## Abstract

Evolution of sex chromosome dosage compensation in mammals remains poorly understood. Ohno’s hypothesis state that evolution of dosage compensation occurred through two steps: first, to compensate the dosage imbalance created due to the degradation of Y chromosome in male, upregulation of X-chromosome happened. Subsequently, transmission of X-chromosome upregulation (XCU) in female led to the evolution of X-chromosome inactivation (XCI) to counteract extra dosage of X-linked genes in female cells. Here, we have profiled gene-wise dynamics of XCU in pre-gastrulation mouse embryos at single cell level and found that XCU is dynamically linked with XCI, however, XCU is not global or chromosome-wide like XCI. Therefore, our result raises question whether XCU driven the evolution of XCI. If so, then why XCI is chromosome wide while XCU is not. We propose that XCI might have evolved independent of XCU and therefore refining the current model is necessary. Separately, we show that higher occupancy of different activating factors at upregulated X-linked genes leads to enhanced transcriptional burst frequency and thereby leads to upregulation. On the other hand, our analysis indicates that extent of upregulation, enrichment of different activating marks differs between ancestral and newly acquired X-linked genes. Altogether, our study provides significant insight into the dynamics and mechanistic basis of evolution of sex chromosome dosage compensation.

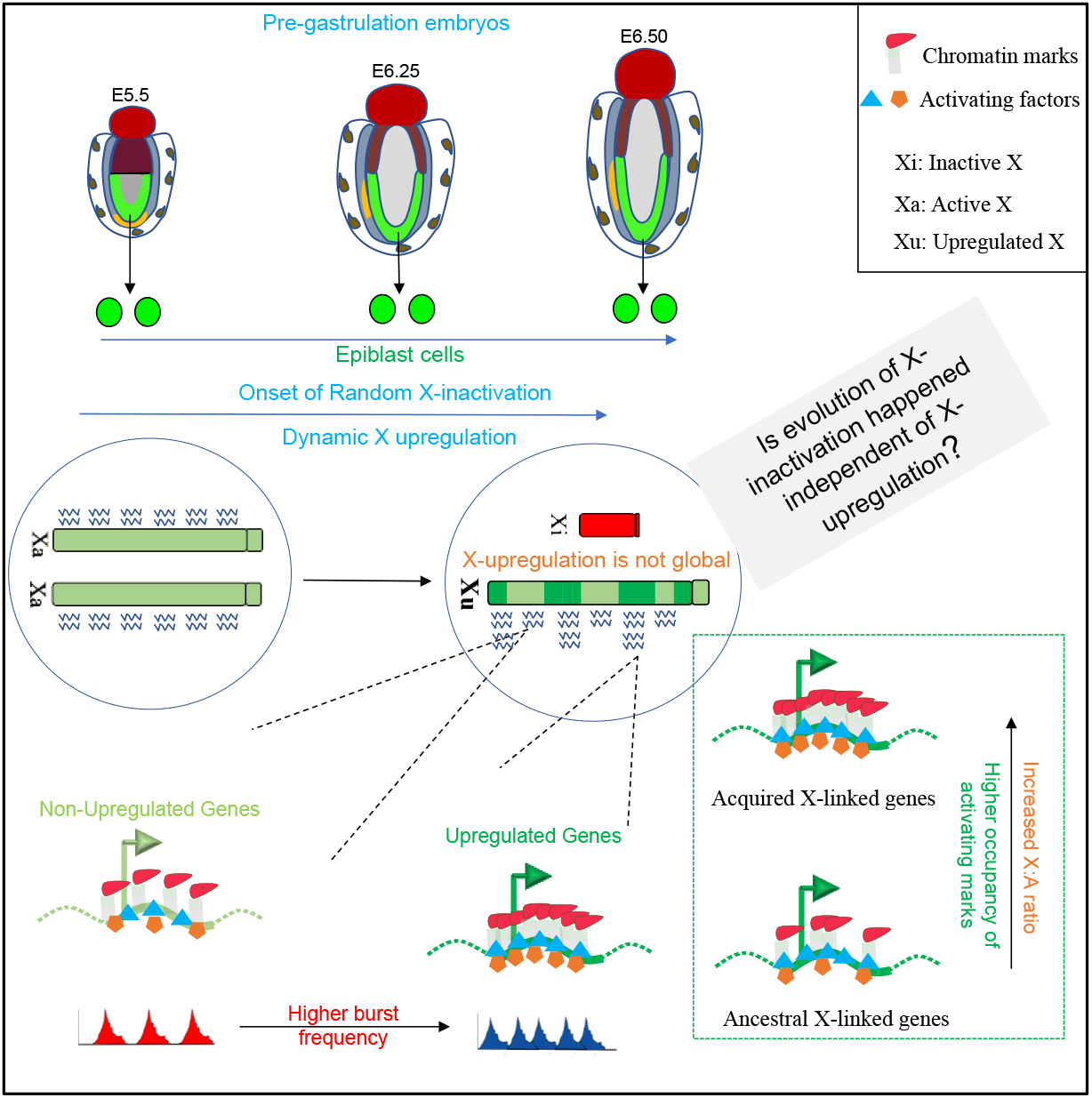

## Introduction

In therian mammals, sex is determined by the sex chromosomes: XX in female and XY in male. The X and Y evolved from a pair of autosomal homologs around 166-180 million years ago through the acquisition of male determining *Sry* gene on one of the chromosomes. During evolution, acquisition of male genes on the Y led to loss of recombination resulting progressive degradation of Y chromosome. Degradation of Y created dosage imbalance between X and autosomal genes in males and between the sexes^1^. In 1967, Ohno hypothesized that evolution of dosage compensation happened through two steps: X-chromosome in male cells was upregulated to two-fold to correct the dosage imbalance related to Y degradation^2^. Subsequently, this X-chromosome upregulation (XCU) was inherited in females and thereby introduced an extra dosage of the X chromosome in female cells. Therefore, to restore optimal dosage from X-chromosome in female cells, the evolution of X-chromosome inactivation (XCI) happened, a process that silent one of the X-chromosome in female mammals ^3^. However, Ohno’s hypothesis was not accepted well for a long time due to the lack of proper experimental evidence. The first evidence of X-upregulation came through the studies based on microarray analysis; however, subsequently, it was challenged through RNA-seq based analysis ^4-9^. Down the line, several independent studies came out both in supporting and refuting Ohno’s hypothesis ^10-19^. However, several recent studies including us have shown that there is indeed presence of upregulated active-X chromosome (X_2a_) in different cell types *in vivo* and *in vitro* ^20^. Nevertheless, whether XCU driven the evolution of XCI or XCI evolved independent of XCU remains an open question.

On the other hand, the extent of XCU, i.e., whether XCU is chromosome wide like XCI or restricted to specific genes, remains poorly understood. Many studies implicated that XCU might be global, though direct evidence showing gene-wise dynamics of upregulation is not well understood ^11,14^. In contrary, several studies implicated that XCU affects dosage-sensitive genes such as components of macromolecular complexes, signal transduction pathways or encoding for transcription factors ^21,22^. To get better insight about this, here we have explored gene-wise dynamics of XCU in pre-gastrulation mouse embryos at single cell level through allele-specific single cell RNA-seq analysis. We found that XCU is neither global nor restricted to dosage sensitive genes only. Next, we have explored what is the mechanistic basis of why some genes are upregulated while others are not from the same active X-chromosome.

On the other hand, we have investigated if XCU is related to the evolutionary timeline of X-linked genes. Based on the evolutionary time point X-linked genes can be categorized into two classes: ancestral X-linked genes, which were present originally from the beginning of sex chromosome evolution and newly acquired X-linked genes. Whether X-chromosome upregulation happens to the newly acquired X-linked genes or restricted to mostly in ancestral genes is another intriguing question. Therefore, we have analyzed the XCU kinetics in ancestral vs. newly acquire X-linked genes to understand the evolution of XCU.

## Results

### Dynamic active-X upregulation upon random XCI in epiblast cells of pre-gastrulation embryos

In female pre-gastrulation mouse embryos, while extraembryonic cells harbor imprinted inactive X, epiblast cells are at the onset of random XCI ^23-27^. We investigated status of XCU in these different lineages of pre-gastrulation embryos through performing allelic/non-allelic gene expression analysis using the available scRNA-seq dataset of E5.5, E6.25, and E6.5 hybrid mouse embryos ^28^ (Fig. 1A).

**Figure 1:**
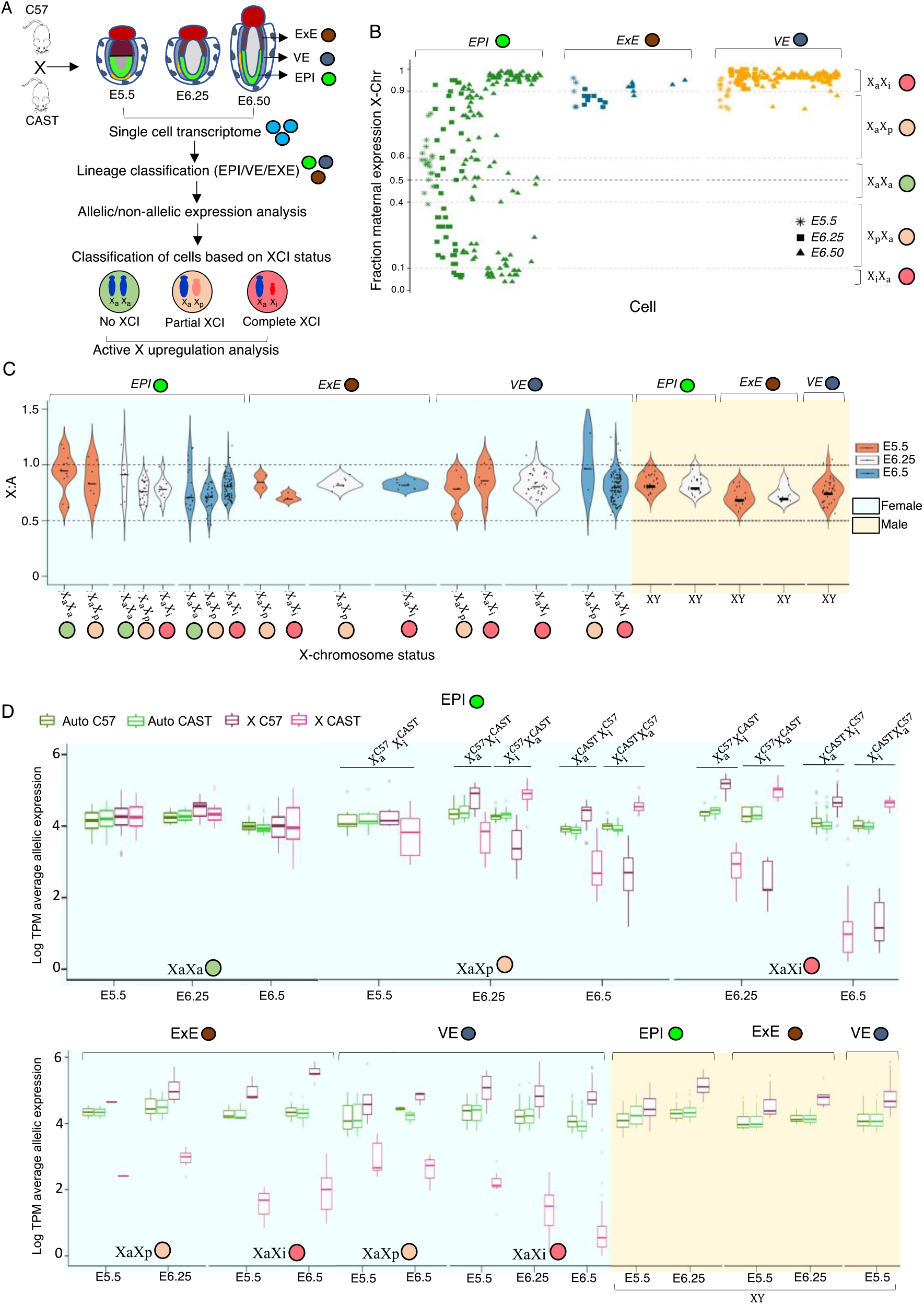
Dynamic X-chromosome upregulation in different lineages of pre-gastrulation embryos. (A) Schematic outline of the workflow: profiling of active X-upregulation in different lineages (EPI: Epiblast, ExE: Extraembryonic ectoderm, and VE: Visceral endoderm) of pre-gastrulation hybrid mouse embryos (E5.5, E6.25, and E6.50) at the single-cell level using scRNA-seq dataset. Hybrid mouse embryos were obtained from crossing between two divergent mouse strains C57 and CAST. (B) Classification of cells based on XCI state through profiling of fraction maternal expression of X-linked genes in the single cells of different lineages of female pre-gastrulation embryos (EPI, ExE, and VE). Ranges of fraction of maternal expression was considered for different category cells: X_a_X_a_ = 0.4-0.6, X_a_X_p_= 0.6-0.9/ 0.1-0.4 and X_a_X_i_ = 0.9-1/0-0.1. (C) X:A ratios represented as violin plots for the different lineages of female (XaXa, XaXp, XaXi) and male cells of pregastrulation embryos. (D) Allelic expression levels of X-linked and autosomal genes in different lineages of female (XaXa, XaXp, XaXi) and male cells of pre-gastrulation embryos.

These embryos were derived from two divergent mouse strains (C57BL/6J and CAST/EiJ) and therefore harbored polymorphic sites across the genome, allowing us to profile gene expression with allelic resolution (Fig. 1A). First, we segregated the cells of E5.5, E6.25, and E6.5 mouse embryos into the three lineages: epiblast (EPI), extraembryonic ectoderm (ExE), and visceral endoderm (VE) based on lineage-specific marker gene expression by t-distributed stochastic neighbour embedding (t-SNE) analysis as described earlier^29^ (Fig. 1A). Next, we categorized the cells of different lineages of the female embryos based on their XCI status: cells with no XCI (X_a_X_a_), partial or undergoing XCI (X_a_X_p_), and complete XCI (X_a_X_i_) through profiling the fraction of maternal allele expression (Fig. 1B). We found that in the EPI lineage lot of cells belongs to X_a_X_p_/X_a_X_a_ category indicating these cells are onset of random X-inactivation, whereas cells of VE/ExE lineage mostly belong to X_a_X_i_ category indicating establishment of imprinted X-inactivation (Fig. 1B). As expected, autosomal genes showed almost equivalent paternal/maternal allele expression, thus validating our allele-specific analysis (Fig S1A). Next, to check the upregulation dynamics, we profiled X:A ratio in the individual cells of different stages/lineages of embryos. If a diploid female cell (X_a_X_i_) has upregulated active X, the X:A ratio should be more than 0.5 and closer to 1. We found that X:A ratio of X_a_X_p_/X_a_X_i_ cells is always > 0.5 and close to 1 despite XCI, indicating the presence of dynamic X-upregulation from the active X chromosome in X_a_X_p_/X_a_X_i_ cells (Fig. 1C). Similarly, male cells had X:A ratio > 0.5 and close to 1 suggesting presence of upregulated X-chromosome (Fig. 1C). For X:A analysis, we made sure that there are no significant differences between the X-linked and autosomal gene expression distribution (Method; Supplementary file 1). Next, to validate the presence of XCU in direct way, we compared the allelic expression pattern from autosome and X-chromosome in individual cells of each lineage. Indeed, we found that the active X expression is always significantly higher than the autosomal allelic expression in X_a_X_p_/X_a_X_i_ cells, corroborating upregulation of gene expression from the active X-chromosome (Fig. 1D). On the other hand, there were no significant differences in active Xs and autosomal allelic expression in X_a_X_a_ cells of epiblast lineage suggesting no upregulation of X-chromosome in absence of XCI (Fig. 1D). As expected, X-chromosome in male cells also showed higher expression than each allele of autosomes (Fig. 1D, Fig S1B,C). Altogether, these analyses suggested (a) dynamic active XCU upon initiation of random XCI in female epiblast cells (b) presence of upregulated X-Chr. in female VE and PE cells with imprinted X inactivation and (c) presence of upregulated X-Chr. in different lineages of male cells of pre-gastrulation embryos as well.

### X-chromosome upregulation is not global

Our X:A analysis in female pre-gastrulation embryos revealed that the X:A ratio of XaXi cells is always lower compared to the XaXa cells (Fig. 1C). This data hinted that X-upregulation of X-linked genes in XaXi cells is partial or all genes do not undergo upregulation. To explore this further, we investigated whether X-chromosome upregulation occurs globally or restricted to certain class of genes. To test this, we profiled gene-wise dynamics of XCU by comparing the expression of X-linked genes from the active X chromosome of X_a_X_i_ cells with the same active allele of X_a_X_a_ cells in EPI E6.5 (Fig. 2A). If active allele of a gene is upregulated in X_a_X_i_ cells, it will show increased expression from the active allele in X_a_X_i_ cells compared to the same active allele of X_a_X_a_ cells. We found that while many X-linked genes showed increased expression from the active-X allele in X_a_X_i_ cells compared to the X_a_X_a_, there were significant number of genes which did not show such increased expression suggesting that all genes do not undergo upregulation (Fig. 2A). Moreover, while some genes showed robust upregulation from the active-X allele, the others were moderately upregulated. Altogether, this result suggested that X-upregulation is not global or do not occur chromosome-wide. On the other hand, surprisingly we found that an adequate number of upregulated X-linked genes showed allele-specificity as they showed upregulation from either C57 or CAST as an active-X suggesting regulation of upregulation of active allele may occur in parent of origin specific manner (Fig. 2B).

**Figure 2:**
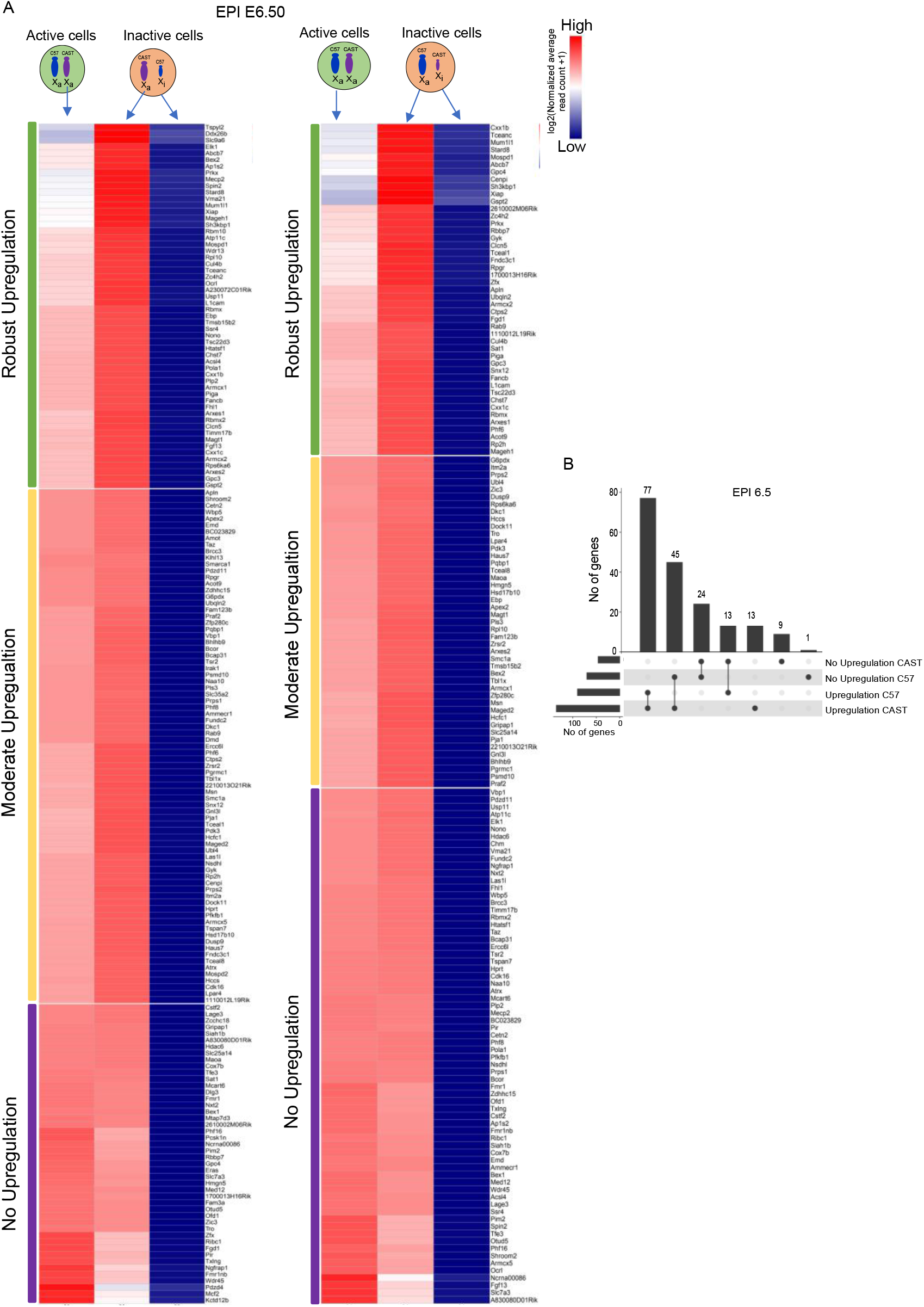
X-chromosome upregulation is not global. (A) Comparison of expression (log normalized reads) of individual X-linked genes from the active allele between the XaXa and XaXi EPI cells of E6.50 embryos (B) Plot representing intersections of allele-wise upregulation of X-linked genes in E6.5 EPI cells.

### Higher occupancy of different activating factors at upregulated X-linked genes versus non-upregulated X-linked genes

Next, we investigated the mechanistic basis of why some genes undergo upregulation and others are not from same active X. For this purpose, we considered the X-linked genes (in E6.5 EPI) having upregulation from both alleles as upregulated genes whereas genes sowing no upregulation from either of the allele categorized as non-upregulated (Fig. 3A). Previous reports stated that XCU is restricted only to dosage-sensitive genes such as genes encoding for large protein complexes, transcription factors, proteins involve in signal transduction etc. Therefore, we analyzed if the upregulated genes in EPI E6.5 is mostly belonging such dosage sensitive genes or not. We identified dosage sensitive X-linked genes through different gene function databases as described in methods. We found that while there are some upregulated genes fall into dosage sensitive category, many genes are not (Supplementary file 2). Moreover, we found that there are many dosage sensitive X-linked genes among the non-upregulated genes. Altogether, there are no significant differences of distribution of dosage sensitive / insensitive genes among the upregulated or non-upregulated genes suggesting XCU is not restricted to dosage sensitive genes only (Supplementary file 2). Additionally, we did not find any relevance of the location of the genes on the X-chromosome or even location of the genes in topologically associated domains (TADs) with the upregulation (data not shown). Next, we asked if there any differences of expression level between upregulated vs non-upregulated genes and found no significant differences (Fig. 3B). Previous studies suggested that X-upregulation is associated with enhanced transcriptional burst frequency. To explore this, we profiled the allelic transcriptional burst kinetics between the upregulated and non- upregulated X-linked genes in the X_a_X_i_ cells of E6.5 EPI. Interestingly, the upregulated X-linked genes showed significantly higher burst frequency compared to the not upregulated X-linked genes (Fig. 3C). However, burst sizes were not significantly different between the two categories of X-linked genes (Fig. 3C). Owing to the higher transcriptional burst frequency of the upregulated genes, next we investigated if upregulated genes have higher occupancy of different transcriptional activating factors compare to the non-upregulated genes. To explore this, we profiled the occupancy of RNA PolII-S5P, RNA PolII-S2P, active chromatin marks such as H3K4me3, H3K36me3 at the transcriptional start site (TSS) and gene body of active allele of upregulated and non-upregulated genes loci through allele-specific Chip-Seq analysis of hybrid female mouse embryonic fibroblast (MEF) cells (Fig 3D). These MEF cells harbor skewed inactive X-chromosome (129S1) and therefore allowed us to differentiate the enrichment of active marks between active and inactive-X through allele-specific analysis. Indeed, we found that while active allele showed significant enrichment of these different active marks, inactive allele had almost no such enrichment thus validating our allele-specific Chip-Seq analysis (Fig 3D). Interestingly, we found that the H3K4me3, RNA PolII (S5P/S2P) showed higher occupancy around the TSS, and gene body regions of the upregulated X-linked genes compared to the non-upregulated genes (Fig. 3D). However, no upregulated gene specific enrichment was observed for H3K36me3. Altogether, these data suggested that enhanced occupancy of different activating marks at the upregulated genes loci might leads to the higher transcriptional burst frequency and thereby leads to the upregulation of X-linked genes.

**Figure 3:**
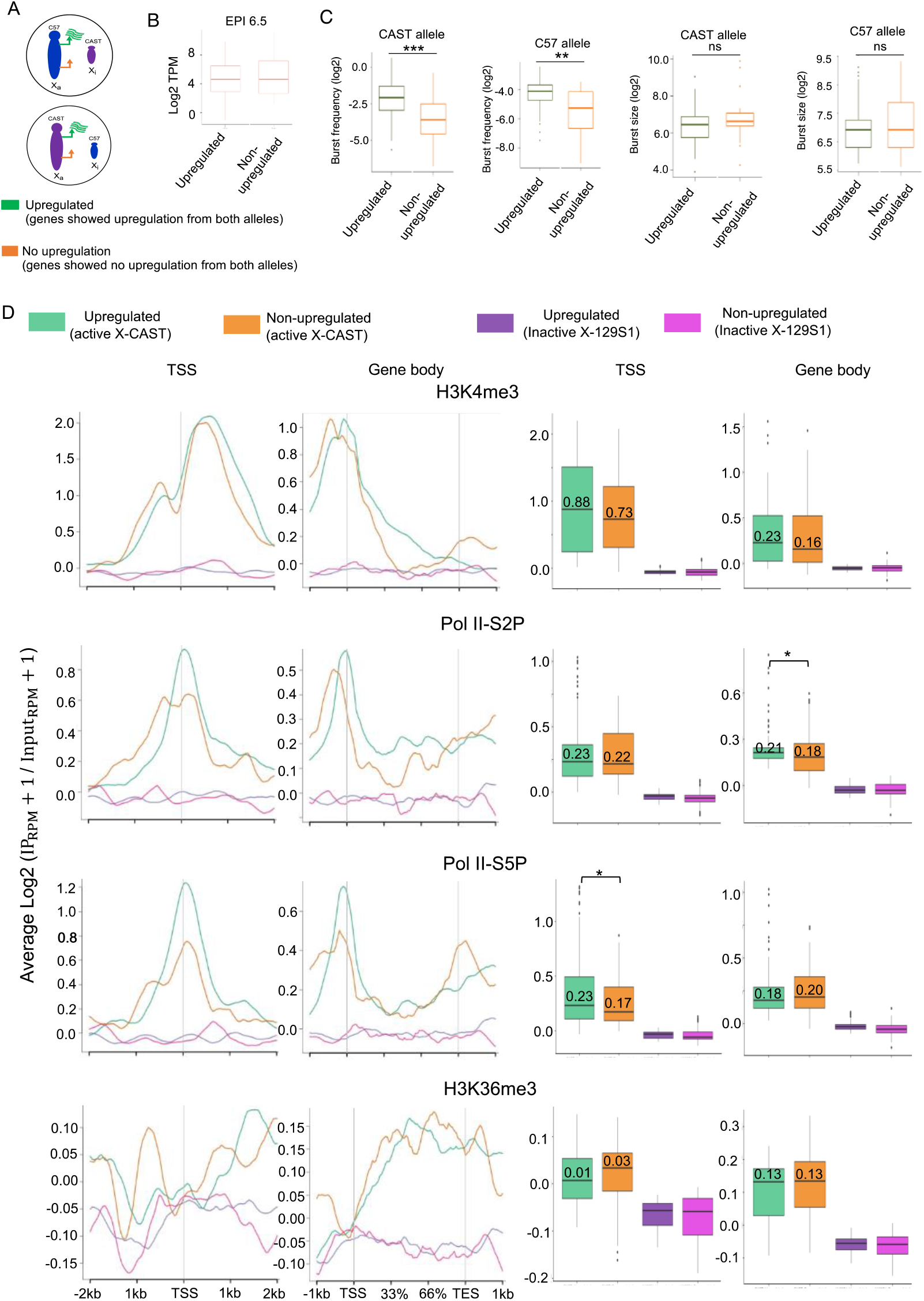
Enrichment of activating factors and increased transcriptional burst frequency leads upregulation of X linked genes. (A) Schematic representation of two category of genes; Upregulated: X-linked genes showing upregulation from both C57 and CAST allele, and non-upregulated: X-linked genes with no upregulation from either of the allele. (B) Comparison of expression levels (log2 TPM) between upregulated and non-upregulated X-linked genes. (C) Comparison of allelic burst frequency and burst size of the upregulated vs non-upregulated X-linked genes in the E6.5 EPI cells (Wilcoxon rank test: *p-value < 0.01*; ** and *p-value < 0.001*; ***). (D) Quantitative enrichment analysis of different activating marks (RNA PolII S5P/S2P, H3K4me3, H3K36me3) at TSS and gene body of upregulated and non-upregulated genes in female MEF cells.

### In-silico model predicts that recruitment probabilities of activating factors as well as availability of activating factors are keys to the upregulation of X-linked genes

To better understand the mechanisms behind XCU through quantitative manner, we hypothesized that the difference in the response to X-upregulation for different X-linked genes could be caused by a difference in the recruitment rates of activation factors of different genes.

To test the validity of the hypothesis, we created an *in silico* model of the X chromosome consisting of two classes of genes, upregulated and non-upregulated. The first category of genes has relatively higher probability of recruitment of activation factors (see Methods). Based on the simulation, we found that the burst frequency, calculated as the fraction of times a gene turns on, is seen to be higher for the upregulated genes as compared to non-upregulated genes for XaXi cells (Fig 4A), thus recapitulating our experimental observations in Fig. 3C. In contrast, simulation in XaXa cells showed no difference between the two classes of genes (Fig 4A). Additionally, we found that this trend of expression level differences holds across the expression matrices generated using the simulations (Fig 4B). Altogether, this analysis suggested that a difference in the range of recruitment probabilities is sufficient to bring about a difference in mean burst frequencies of upregulated and non-upregulated genes.

**Figure 4:**
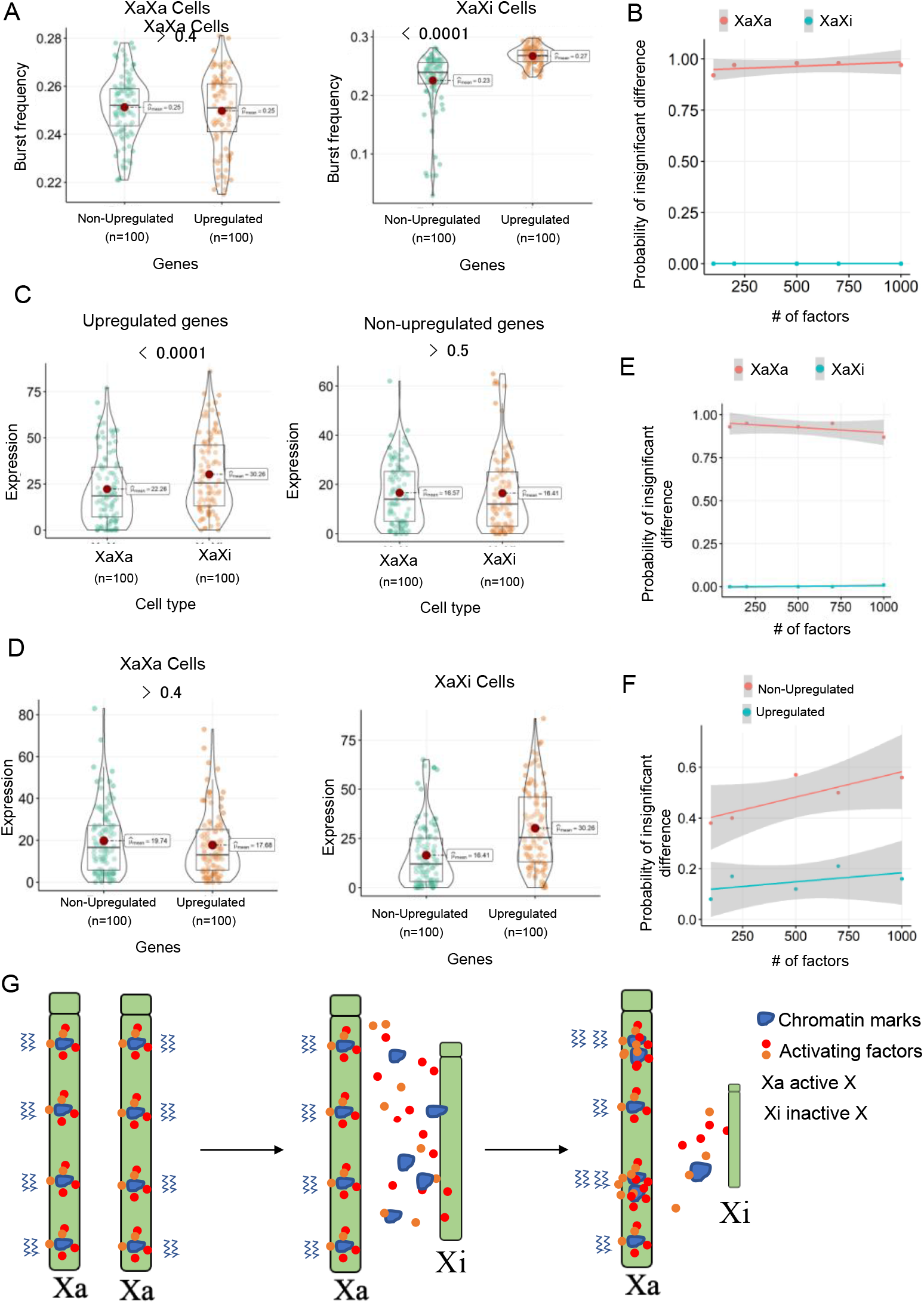
In silico model explains a possible mechanism for XCU. (A) Burst frequency distributions for upregulated and non-upregulated genes in XaXa and XaXi cells from the model. (B) Fraction of cells having insignificant difference between the burst frequency of upregulated and downregulated genes for XaXa cells (red), XaXi cells (blue). (C) Expression level distribution for upregulated and non-upregulated genes from active allele from XaXi (orange) and an allele from XaXa (green) in the model (D) Difference in the expression levels of upregulated vs non-upregulated genes of XaXa and XaXi cells. (E) Same as B but for difference in expression levels of up and non-upregulated genes. (F) Fraction of in-silico cells that show insignificant difference in the expression levels of upregulated genes of XaXa and XaXi (red) and non-upregulated genes between XaXa and XaXi (blue). (G) Model representing that increase of activating factors availability upon XCI and followed by enrichment of these activating factors at the loci of genes with higher recruitment probability on the active X chromosome leads to the upregulation of those X-linked genes.

We then compared the expression levels of upregulated and non-upregulated genes across XaXa and XaXi cells. The difference in expression levels is statistically significant for the upregulated genes but not so for non-upregulated genes (Fig 4C), suggesting that a difference in recruitment probabilities can also bring about gene upregulation in the active chromosome upon XCI. This result is also supported by the fact that within the two kinds of cells (XaXa and XaXi), the difference between upregulated and non-upregulated genes could be seen only for XaXi cells, where there was a difference in the recruitment probabilities (Fig 4D-E). We then examined whether the activation factors available to chromosome play any role in determining this difference. We generated expression matrices for different levels of activation factors and found that at lower levels of activation factors, the upregulation is quite noisy, because the downregulated genes also show significant difference in expression between XaXa and XaXi for more than 60% cells (Fig 4F). As the number of factors increases, the percentage of downregulated genes showing upregulation decreases, making upregulation less stochastic (Fig 4F).

### Extent of upregulation and occupancy of different activating marks differs between ancestral and newly acquired X-linked genes

Next, we investigated whether there is evolutionary constraint to the X-upregulation. Based on the evolutionary time point X-linked genes can be categorized into two classes: ancestral X-linked genes, which were present originally from the beginning of sex chromosome evolution and newly acquired X-linked genes. To test if upregulated genes are restricted to one of these evolutionary X linked gene classes, we collated the upregulated and non-upregulated genes with their orthologs in chicken (ancestral X-linked genes) and found that most of the upregulated genes belong to ancestral class of X-linked genes, however, non-upregulated genes had equivalent distribution to ancestral and acquired X-linked genes (Fig. 5A). Based on this preliminary observation, we hypothesized that X upregulation might be mostly confined to the ancestral class of X-linked genes. To test this further, we profiled X:A ratio for ancestral and acquired X-linked genes in XaXi cells of female E6.5 EPI and VE. Surprisingly, we found that newly acquired X-linked genes showed significantly higher X:A ratio compared to the ancestral genes (Fig. 5B). This was consistent in male cells as well (Fig. 5B). Based on this, we concluded that extent of X-upregulation is higher for acquired X-linked genes compared to the ancestral genes. Next, to find out the mechanisms behind this we profiled transcriptional burst kinetics and occupancy of different activating marks in ancestral and acquired X-linked genes. We profiled allelic burst frequency and burst size in E6.5 EPI XaXi cells. Our analysis revealed that there were no significant differences of allelic transcriptional burst frequency or burst size between acquired and ancestral X-linked genes. However, both classes of genes (ancestral and acquired) had significantly higher transcriptional burst frequency compared to the autosomal genes corroborating the fact that both classes undergo upregulation (Fig. 5C). Surprisingly, we found that the burst size of ancestral and acquired genes were significantly low compared to the autosomal genes (Fig. 5C). On the other hand, to explore further if there any differences in occupancy of different active marks between acquired and ancestral X-linked genes we profiled the enrichment of different marks around the TSS, and gene body regions of ancestral and acquired X and autosomal genes (Fig. 5D). We profiled the occupancy of RNA PolII-S5P/S2P, H3K4me3, H3K36me3 in hybrid female MEF cells using the available Chip-seq data, same datasets used for analysis in Fig.3. We found that inactive X-chromosome (129S1) had almost no enrichment for these marks as revealed by the allelic enrichment analysis (Fig. S2). Considering this fact, we profiled enrichments of these marks at ancestral and acquired loci through non-allelic analysis to have the better representation of both classes of genes (ancestral and acquired) and thereby to have more accurate picture of enrichment of different activating marks as through allelic analysis we lose many genes and reads due to the lack of SNPs. For autosomal genes, we found that enrichment of marks is almost equal between the two allele (Fig. S2) and therefore we divided total enrichment score by 2 to have enrichment corresponding to one allele. Moreover, we eliminated potential escape genes from our analysis as they will have enrichment of these marks from inactive allele as well. On the other hand, to avoid any bias created due to the low expressed genes we eliminated low expressed genes (≤ 0.5FPKM) as well as multicopy genes from our analysis. We found that acquired X-linked genes showed significantly higher occupancy for all the marks except H3K4me3 around the TSS and gene body regions compared to the ancestral X-linked genes (Fig. 5D). Separately, we also profiled enrichment of H4K16ac in male MEF cells and found that acquired X-linked genes have significantly higher occupancy around the TSS, and gene body regions compared to the ancestral X-linked genes (Fig. 5D). Altogether, these data suggested that acquired X-linked genes have higher occupancy of different activating marks compared to the ancestral X-linked genes. On the other hand, we did not observe such trends between ancestral and acquired autosomal genes suggesting this is restricted to the evolution of X-chromosome (Fig 5D). Moreover, we found that ancestral X-linked genes had significant higher enrichment of different marks (H3k4me3, RNA PolII S5P/2P) at TSS compared to the autosomal ancestral genes (Fig 5D). Similarly, acquired X-linked genes showed significant higher enrichment of different marks at TSS (H3k4me3, RNA PolII S5P/2P and H4k16ac) as well as in gene body (RNA PolII S5P/2P and H4k16ac) compared to the autosomal acquired genes suggesting both of the classes undergo X-upregulation.

**Figure 5:**
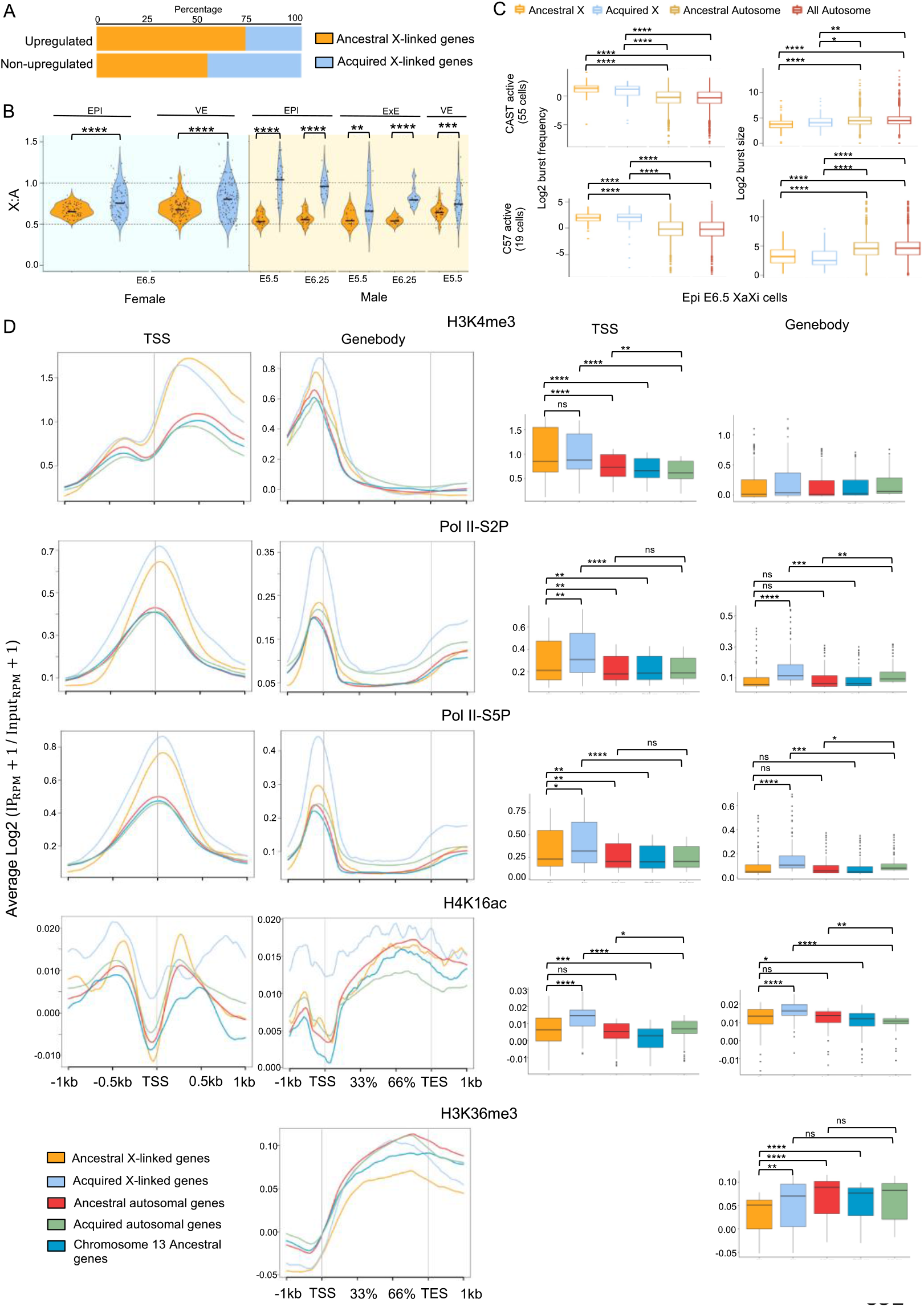
X-upregulation is mostly confined to ancestral X-linked genes. (A) Plot representing the percent of upregulated and non-upregulated X-linked genes belonging to ancestral or acquired X-linked gene category. (B) X:A ratio analysis for ancestral and acquired X-linked genes in female (XaXi) and male cells of EPI, EXE and VE. (C) Comparison of allelic burst frequency and burst size between ancestral or acquired X-linked genes in EPI 6.5 XaXi cells. (D) Comparison of enrichment of different activating marks (RNA PolII S5P/S2P, H3K4me3, H4K16ac and H3K36me3) at TSS and gene body of ancestral vs acquired X-linked and autosomal genes.

**Figure 6:**
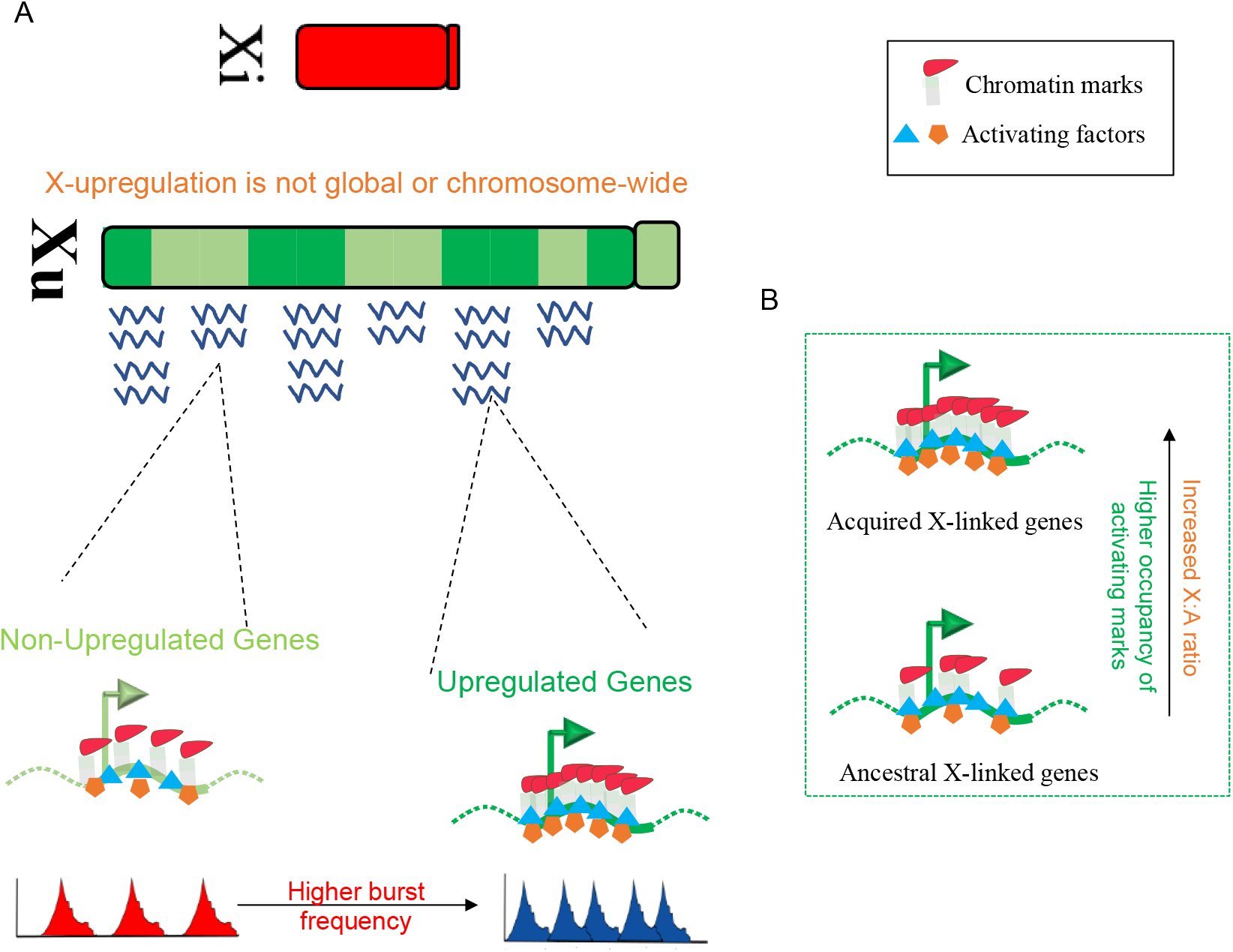
Model representing X-upregulation dynamics and mechanisms. (A) X-upregulation is not global or chromosome-wide. Upregulated X-linked genes show higher occupancy of different active marks and have higher transcriptional burst frequency compared to the non-upregulated genes. (B) Extent of X-upregulation, occupancy of different activating marks differs between ancestral and newly acquired X-linked genes.

## Discussion

Here, we have profiled the XCU dynamics at single cell level in female pre-gastrulation mouse embryos. We found that in EPI cells (E5.5, E6.25, E6.5), which are at the onset of random XCI, XCU is dynamically linked with the XCI. Moreover, VE and ExE cells, which undergo imprinted XCI also showed presence of upregulated active-X chromosome. Altogether, these results suggested that two X-chromosomes expression state is highly plastic in nature towards balancing the optimal X chromosome dosage during pre-gastrulation. Our result is consistent with a recent report by Lentini et al.^30^. However, we found that though XCU is dynamically linked to XCI, XCU is not global like XCI. Analysis of gene-wise dynamics of XCU revealed that many X-linked genes do not undergo upregulation (Fig 2A). This was also reflected in X:A ratio analysis, where we found that X:A ratio of XaXi cells tends to be lower compared to the XaXa cells (Fig. 1C) Therefore, our result raises question about the current theory of evolution of XCI that the evolution of XCI happened to counteract the upregulation of X in female cells as proposed by Ohno. If so, then why XCI is global and occurs to the majority of X-linked genes while XCU is not. Based on this, we propose that XCI might have evolved independent of XCU and in the future more investigation is necessary to test whether XCI evolved from a completely different perspective. It also may be possible that initially XCI evolved to the genes undergoing upregulation and later it became a global phenomenon for some unknown reasons. An alternative model (parental antagonism model) of evolution of XCI was proposed by Haig suggesting that XCI may have evolved initially in the form of genomic imprinting related to parental conflicts ^22,31,32^. Haig’s hypothesis also predicts that imprinted paternal XCI rather than the random XCI to be the ancient form of XCI. Indeed, in marsupials and in some tissues of placentals XCI is always occurs on paternal X. However, further investigation is necessary to validate the Haig’s model of evolution of XCI. Separately, XCI might be very old as it is present in marsupials as well. In the future, it would be interesting to examine whether XCU is also present in marsupials or not which may clear who evolved first XCI or XCU. On the other hand, previous studies implicated that XCU affects dosage-sensitive genes such as components of macromolecular complexes, signal transduction pathways or encoding for transcription factors ^21,22^. However, we found that upregulated genes are not restricted to dosage sensitive genes only and there are many dosage sensitive genes, which do not undergo upregulation as well (Supplementary file 2).

Next, our analysis unveils the mechanistic basis of how some genes get upregulated and others are not. Previous studies have reported that the X linked genes from active X-chromosome have greater enrichment of the active chromatin marks compared to the autosomal genes ^14^. We found that upregulated X-linked genes have the higher occupancy of different transcriptional activating marks such as H3K4me3, RNA PolII S2P (gene body), RNA PolII S5P (TSS) compared to the non-upregulated genes (Fig. 3D). Interestingly, transcriptional burst frequencies were significantly enhanced in upregulated genes compared to the non-upregulated genes (Fig. 3C). Taken together, our results unveils that the enrichment of activating marks might leads to higher transcriptional burst frequencies and thereby results upregulation of X-linked genes. Additionally, our in-silico analysis shows that a difference in the range of recruitment probabilities of different activating factors is sufficient to bring about a difference in the mean burst frequencies of upregulated and non-upregulated genes (Fig. 4). Importantly, availability of the activation factors also play important role in determining these differences. We predict that upon XCI numerous trans-acting factors leave the inactivating-X and thereby leads to global increase of the number of activating factors. We show that upon such increase of activating factors availability, genes with higher recruitment probability gets enriched with these factors and thereby brings upregulation of those genes (Fig 4G).

Finally, we investigated the evolutionary constraint behind X-upregulation. We found that both ancestral and newly acquired X-linked genes undergo XCU, however, extent of X-upregulation differs between ancestral and newly acquired X-linked genes. We found that X:A ratio for acquired genes is significantly higher compared to the ancestral genes (Fig 5B). Similarly, enrichment analysis of different active marks revealed that both ancestral and newly acquired X-linked genes had higher enrichment of different marks (TSS: H3k4me3, RNA PolII S5P/2P, H4k16ac; Gene body: H4k16ac) compared to the autosomal ancestral and acquired genes (Fig 5D). However, in most cases acquired X-linked genes showed higher enrichment of different marks compared to the ancestral X-linked genes (Fig 5B). Altogether, these data suggested that extent of upregulation for acquired X-linked genes is higher compared to the ancestral genes. However, in terms of burst frequency, we did not observe significant differences (Fig 5C).

In summary, we show that although the two X-chromosomes’ expression state is highly plastic in nature towards balancing the optimal X chromosome dosage during development, XCU is not global like XCI. Therefore, our result raises question why XCI is chromosome wide while XCU is not if evolution of XCI happened to counteract the upregulation of X in female cells as proposed by Ohno. Based on this, we propose that evolution of XCI might have happened independent of XCU and therefore refining the current model is urgently needed. Importantly, we found that enhanced occupancy of different activating marks at the upregulated genes loci might leads to the higher transcriptional burst frequency and thereby leads to the upregulation. Finally, our analysis revealed that extent of X-upregulation differs between ancestral and newly acquired X-linked genes.

## Methods

### Data acquisition

Pre-gastrulation embryo single-cell RNA-seq dataset for E5.5 and E6.25 generated from hybrid mouse embryos (C57BL/6J × CAST/EiJ) and E6.5 from hybrid mouse with the reciprocal cross (CAST/EiJ × C57BL/6J) was retrieved under the accession code-GSE109071 ^28^.

### Read alignment and counting

RNA-seq reads were aligned to the mouse reference genome mm10 using STAR ^33^. The reads aligned were then counted using HTSeq-count. To avoid the dropout events due to low amount of mRNA sequenced within single cells, we used a statistical imputation method scImpute^34^, which identifies the likely dropouts without introducing any bias in the rest of the data. Expression levels of transcripts was computed using *Transcripts per million* (TPM) method.

### Sexing of the embryos

For sex-determination of the pre-gastrulation embryos, an embryo was classified as male if the sum of the read count for the Y-linked genes (Usp9y, Uty, Ddx3y, Eif2s3y, Kdm5d, Ube1y1, Zfy2, Zfy1) in each cell of an embryo was greater than 12, rest were considered as female embryos.

### X:A ratio analysis

We calculated the X:A ratio for different lineages of pre-gastrulation embryo by dividing the median expression (TPM) of X-linked genes with the median expression (TPM) of the autosomal genes. For this analysis, we considered those X-linked and autosomal genes having ≥ 0.5 TPM. We also applied an upper TPM threshold which corresponded to the lowest 90 th centile of TPM expression to avoid any differences between the X-linked and autosomal gene expression distribution. Also, statistical Kolmogorov-Smirnov’s test was performed which again validated the non-significant differences in the levels of gene expression distribution between X and autosomal genes. We excluded the escapees of X-inactivation and the genes in the pseudo autosomal region from our analysis.

### Allele-specific expression analysis

We first constructed in silico CAST specific parental genome by incorporating CAST/EiJ specific SNPs (https://www.sanger.ac.uk/science/data/mouse-genomes-project) into the mm10 genome using VCF tools. Reads were mapped onto C57BL/6J (mm10) reference genome and in silico CAST genomes using STAR allowing no multi-mapped reads. To exclude any false positives, we only considered those genes with at least 2 informative SNPs and minimum 3 reads per SNP site. We took an average of SNP-wise reads to have the allelic read counts. After normalization of allelic read counts, we considered only those genes for downstream analysis which had at least 1 average reads across the cells of each lineage from a specific developmental stage for pre-gastrulation embryos. Further, only those single-cells were considered for downstream analysis which showed at least 10 X-linked gene expressions. Allelic ratio was obtained individually for each gene using formula = (Maternal/Paternal reads) ÷ (Maternal reads + Paternal reads).

### Identification of ancestral and multicopy genes

For ancestral gene identification, we used the HGNC database HCOP (https://www.genenames.org/tools/hcop/) tool, which uses different well established inference methods to retrieve the orthologous genes. From the bulk data retrieval (http://ftp.ebi.ac.uk/pub/databases/genenames/hcop/), first we retrieved the human vs mouse orthologs and human vs chicken orthologs and subsequently we merged both ortholog datasets on human genes to identify mouse related orthologs on chicken. We considered only high confident orthologs genes, which were present in >=3 inference methods for our analysis. Multicopy genes for mouse were retrieved from the *Ensembl* biomart database. We considered genes which are having paralog >=90% as multicopy genes as described earlier ^35,36^.

### Transcriptional burst kinetics

We used SCALE and Txburst to profile allelic transcriptional burst kinetics ^37^. For this analysis, we used genes with at least 5 average read counts across the cells of EPI E6.5 considering that single cells are more prone to dropout event. Additionally, escapee genes were removed from our analysis.

### ChIP-seq analysis

To estimate the enrichment of different activating marks across TSS and gene body, we retrieved the available ChIP-seq datasets for MEF cells from GSE33823 ^38^, GSE36905 ^39^ for H3K4me3, H3K36me3, RNAPolII S5P/S2P and GSE97459 ^40^ for H4K16ac. First, we created A ‘N-masked genome in-silico using SNPsplit (0.4.0) ^41^. The reads for all samples are then mapped to this N-masked genome with Bowtie2 (-N 1) ^42^. Duplicate reads were removed with Picard ‘MarkDuplicates’ [‘REMOVE_DUPLICATES=TRUE’] and blacklisted regions were removed according to encode consortium. SNPsplit was then used to create allele-specific BAM files by segregating the aligned reads into two distinct alleles (129S1/SvImJ and CAST/EiJ). Enrichment plots and quantification were created Using ngs.plot (-AL bin-MW 15) ^43^. For filtering genes based on expression (>=0.5 FPKM) for enrichment analysis, we considered RNA seq data of MEF cells from GSE86653 (GSM2308380).

### Identification of dosage sensitive genes

To identify dosage sensitive X-linked genes, we profiled the associated biological function of the genes through Signal related function (http://geneontology.org/) ^44^, Transcription factor databases (http://bioinfo.life.hust.edu.cn/AnimalTFDB/#!/, https://sunlab.cpy.cuhk.edu.hk/mTFkb/) ^45,46^, genes involved in protein complex databases (http://mips.helmholtz-muenchen.de/corum/#) ^47^, dosage sensitive genes (human) database (https://www.clinicalgenome.org/) ^48^, Housekeeping genes databases (https://housekeeping.unicamp.br/, Li et al., 2017) ^49,50^. We called a gene as dosage sensitive if it has association with at least one of the category/databases (Supplementary file 2).

### Simulation

The *in-silico* chromosome: The chromosome is modelled as a set of genes. Each gene has a probability of recruitment of transcriptional activation factors (epigenetic factors, transcription factors etc.). Based on probability, genes were classified into two categories: upregulated and non-upregulated, with decreasing probabilities respectively. We took 100 each of these genes and positioned them at random positions on the chromosome. The recruitment happens from a column of activation factors, with 1000 molecules placed in front of every gene. At every time step, each gene can recruit an activation factor from the corresponding position on the column, depending upon the recruitment probability of the gene. The activation factors are modelled to have a discrete diffusion, in that the reduction in activation factors at any position is filled in by the neighbouring positions. The recruitment onto the gene is modelled as a cooperative process, with the probability of recruitment increasing sigmoidally with increase in the extent of recruitment. Each gene has a probability of activation, also modelled as a sigmoidal function of the extent of recruitment on the gene. The probability of inactivation for each gene is 0.5. An active gene produces one RNA molecule at every time step. These RNA molecules degrade with a probability of 0.35.

Simulation procedure: At each time step, the recruitment of activation factors happens to all genes if the recruitment probability for the genes is higher than a uniformly random number generated. After recruitment, diffusion is carried out to normalize the activation factor column. Then, genes are turned on/off with the on/off probability as described above. All active genes produce one RNA molecule (making the time scale of the simulation the same as transcription). RNA degradation also is implemented one molecule at a time with a probability of 0.35. For sensitivity analysis, we generated 100 *in silico* cells, each represented by a chromosome described above.

### Statistical tests and plots

All statistical tests and plots were performed in R version 3.6.3.

## Author’s Contribution

SG conceptualized and supervised the study. HCN and DC performed data analyses. KH and MKJ worked on mathematical modelling. SG, MKJ, HCN, DC, and KH wrote, edited, and proofread the manuscript.

## Acknowledgments

This study is supported by DBT grant (BT/PR30399/BRB/10/1746/2018), DST-SERB (CRG/2019/003067), DBT-Ramalingaswamy fellowship (BT/RLF/Re-entry/05/2016) and Infosys Young Investigator award to SG. We also thank DST-FIST [SR/FST/LS11-036/2014(C)], UGC-SAP [F.4.13/2018/DRS-III (SAP-II)] and DBT-IISc Partnership Program Phase-II (BT/PR27952-INF/22/212/2018) for infrastructure and financial support.

---

**Figure S1:**
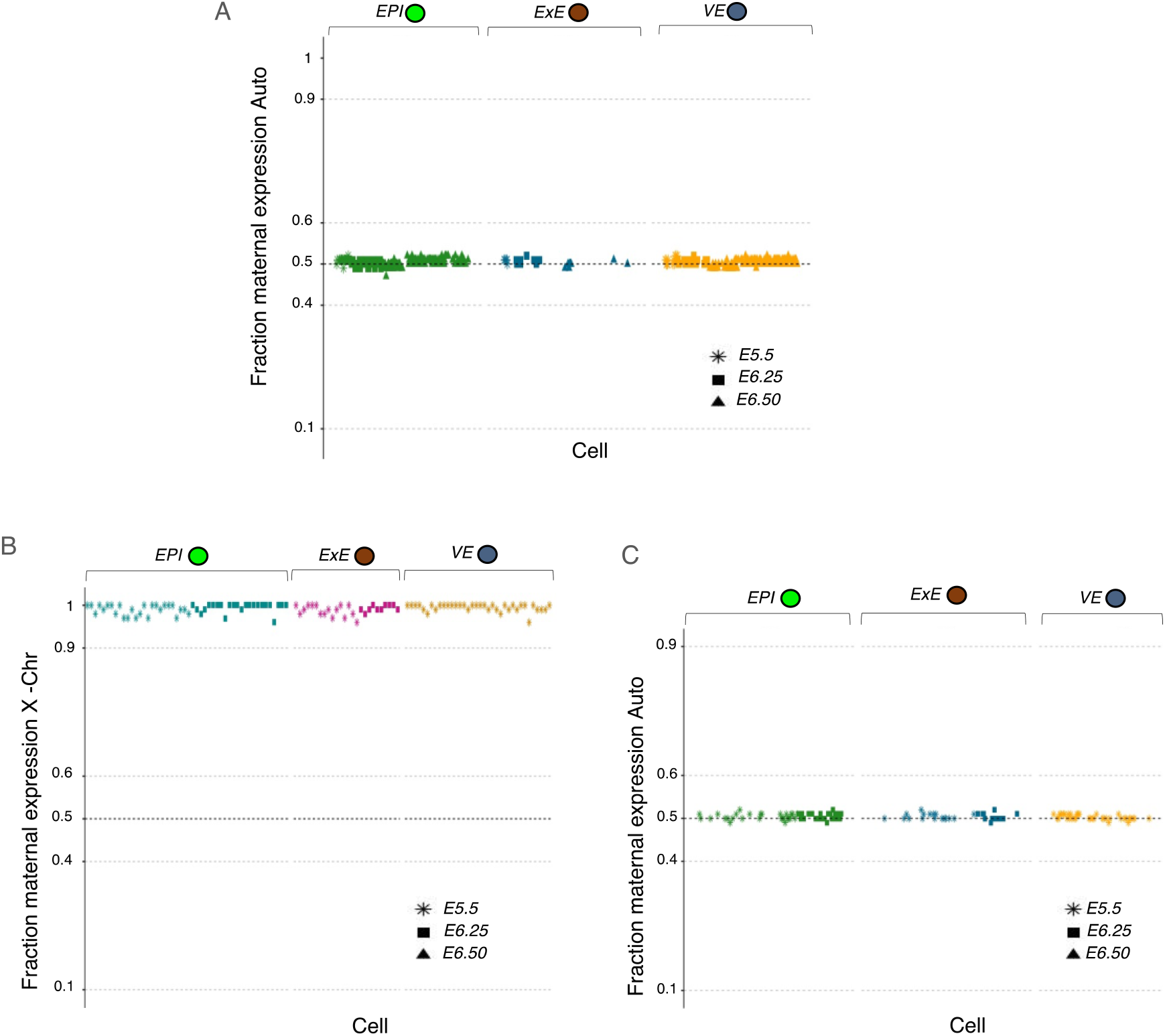
Allelic expression of autosomal genes. (A)Female cells fraction maternal expression of Autosomal genes in the single cells of different lineages of pre-gastrulation embryos (EPI, ExE, and VE). (B) Male cells fraction maternal X-linked genes in single cells of different lineages of pre-gastrulation embryos (EPI, ExE, and VE). (C) Male cells Fraction maternal Autosomal genes in single cells of different lineages of pre-gastrulation embryos (EPI, ExE, and VE).

**Figure S2:**
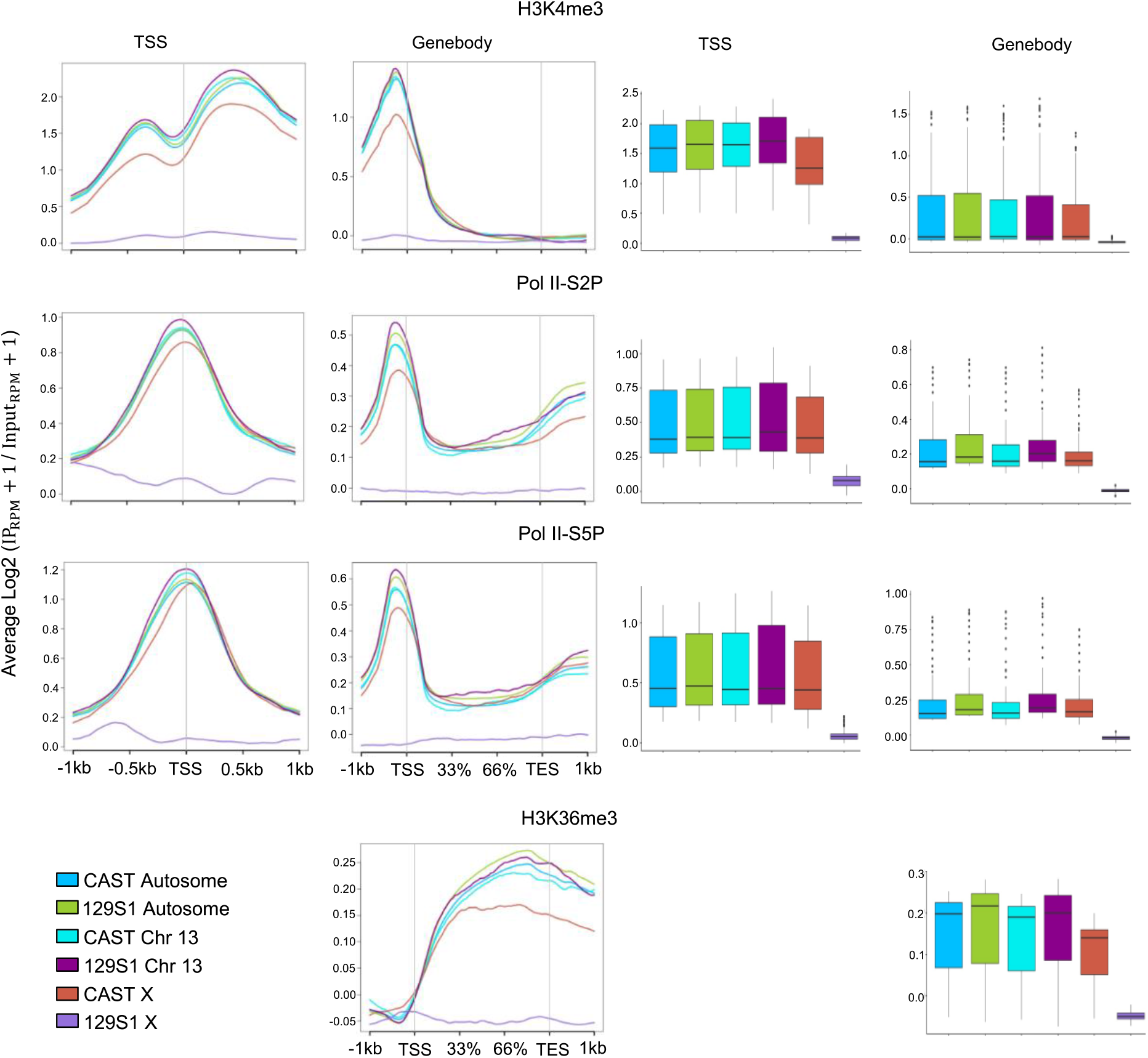
Allele-specific enrichment analysis and quantification of different active marks (RNA PolII S5P/S2P, H3K4me3 and H3K36me3) at TSS and gene body on the X-chromosomes, autosomes and Chr 13.

